# Pathogenic Fusarium verticillioides and Ophiostoma clavatum Associated with Ips acuminatus in Ukraine

**DOI:** 10.1101/2025.11.11.687790

**Authors:** Yurii M. Yusypovych, Yuliia I. Shalovylo, Oleh Y. Kit, Volodymyr O. Kramarets, Volodymyr K. Zaika, Mykola M. Korol, Vasyl V. Lavnyy, Hryhoriy T. Krynytskyy, Valentina A. Kovaleva

## Abstract

Over the past two decades, dieback of *Pinus sylvestris* L. stands has increased across Europe, largely due to mass outbreaks of the bark beetle, in particular, *Ips acuminatus* Gyll. (Coleoptera: Curculionidae). This beetle causes mechanical damage and vectors pathogenic fungi, including ophiostomatoid species that induce blue-stain. *Ophiostoma clavatum* Math.-Käärik is the most frequently reported fungal associate, yet its occurrence had not been documented in Ukraine. While ophiostomatoid fungi are well studied in pine pathogenesis, the role of fast-growing associates such as *Fusarium* spp., remains poorly understood. This study aimed to identify the dominant *Ophiostoma* and *Fusarium* species associated with *I. acuminatus* in western Ukraine and to evaluate their pathogenicity on pine seedlings. Isolates from beetle abdomens and blue-stained wood were identified as *O. clavatum* based on morphology and molecular markers (ITS, TUB, TEF1-α). Pathogenicity tests showed that *O. clavatum* acts as a weak phytopathogen. The dominant *Fusarium* morphotype from blue-stained wood was identified as *Fusarium verticillioides* (Sac) Nirenberg, which induced necrosis and tissue maceration on pine seedlings. In dual culture, *F. verticillioides* displayed strong competitive dominance over *O. clavatum*. This study provides the first record of *O. clavatum* associated with *I. acuminatus* in Ukraine, extending its known European distribution. The observed pathogenicity and competitive ability of *F. verticillioides* suggest it may contribute to Scots pine decline, warranting further investigation.

## 1 INTRODUCTION

Scots pine (*Pinus sylvestris* L.) is a widely distributed forest species of major ecological and economic importance. It is highly adaptable, thriving on nutrient-poor soils and tolerating extreme conditions. However, climate change-induced fluctuations impair its photosynthetic activity and weaken its defenses against stress factors such as bark beetle attacks (Dobbertin et al. 2007; Brichta et al. 2024). *Ips* species, traditionally secondary pests attacking weakened or damaged pines, have recently increased in impact due to extreme weather events like droughts and heatwaves. These conditions reduce forest resilience and alter pest dynamics (Wermelinger et al. 2008; Siitonen 2014). Rising pine damages linked to *Ips acuminatus* Gyll. (Coleoptera: Curculionidae) have been reported across Europe, including France, Switzerland, Italy, Finland, Ukraine, Czechia, and Poland (Lieutier et al. 1991; Dobbertin et al. 2007; Wermelinger et al. 2008; Colombari et al. 2012; 2013; Siitonen 2014; Davydenko et al. 2017; Knížek and Liška, 2023; Jankowiak et al. 2023; Yusypovych et al. 2023). Bark beetles form complex associations with fungi, particularly ophiostomatoid species (blue-stain fungi), which discolor colonized sapwood (Wingfield et al. 1993; Kirisits 2004; Linnakoski et al. 2012). *I. acuminatus* is closely linked to these fungi, with *Ophiostoma clavatum* Math.-Käärik being the most frequently reported associate across multiple studies (Mathiesen, 1950, 1951; Rennerfelt, 1950; Mathiesen-Käärik 1953; Francke-Grosmann 1963; Käärik 1975; Linnakoski et al. 2016).

Since 2010, drought and bark beetle outbreaks have caused extensive dieback of *P. sylvestris* in Ukraine, affecting some 79,000 hectares of pine stands in 2022. The main pests are the aggressive engravers *I. acuminatus* and *I. sexdentatus* (Davydenko et al. 2017, 2021). A geographic variation exists in the fungal communities vectored by *I. acuminatus*. The *O. clavatum* complex, frequently found in Italy and Sweden, was absent from eastern Ukraine; the species present there belonged to the *O. ips*, *O. minus*, and *O. piceae* complexes (Davydenko et al. 2017; de Beer et al. 2022). These regional differences prompted us to study the *Ophiostoma* species associated with *I. acuminatus* in western Ukraine, which has distinct climatic conditions.

In addition to ophiostomatoid fungi, pine bark beetles carry other aggressive pathogens. *Diplodia sapinea* and *Fusarium circinatum* have been found in *I. sexdentatus* galleries, and co-infections with ophiostomatoid fungi may worsen tree decline (Elvira-Recuenco et al. 2020). Notably, *Fusarium* species comprised 3.17% of the *I. acuminatus* microbiome (Chakraborty et al. 2020), suggesting this beetle may vector fusaria.

This study aimed to: (1) determine *Ophiostoma* species associated with *I. acuminatus* in Lviv region (western Ukraine); (2) isolate and characterize the dominant fast-growing fungi, particularly *Fusarium* spp., associated with blue-stain; and (3) assess the pathogenicity of selected *Ophiostoma* and *Fusarium* isolates on pine seedlings.

## 2 MATERIALS AND METHODS

### 2.1 Study site and sampling

This study was conducted in August 2022 in a 50–70-year-old Scots pine stand in the Lviv region, Ukraine (50°23′N, 23°66′E). Within a medium-sized *I. acuminatus*-infested spot (Colombari et al. 2013), five living pine trees showing visible infestation symptoms, including needle discoloration, crown thinning (>40%), and beetle entry holes were selected. The trees were felled, and logs with beetle galleries from upper stem sections and branches were collected and transported to the laboratory on the same day. Adult beetles and larvae were present in the galleries at the time of sampling.

### 2.2 Culture-based isolation

In the laboratory, twenty *I. acuminatus* beetles per tree were collected from galleries using fine forceps. Insect samples were rinsed with sterile distilled water, surface-sterilized in 96% ethanol for 3 minutes, and then rinsed twice with sterile water. The efficacy of the sterilization was verified by plating the final wash onto 2% potato dextrose agar (PDA) and incubating at 24 °C for 5 days. Head-pronotums with wings and legs were aseptically removed. Abdominal segments of beetles from each tree were pooled and transferred to 2-ml microcentrifuge tubes containing 10 mM sterile phosphate-buffered saline (PBS). The samples were homogenized on ice using a plastic pestle, followed by vortexing for 1 minute at 2400 rpm (Biosan, Latvia). The homogenates were centrifuged at 4000 rpm for 5 min to separate the microbial suspension from insect tissue debris (Hu et al. 2015). Supernatants were plated onto 2% PDA supplemented with antibiotics (100 µg/ml ampicillin and 50 µg/ml chloramphenicol), and incubated at 24°C in the dark. All procedures were carried out under sterile conditions using a microbiological safety cabinet (Walker, Glossop, UK). Petri dishes were examined daily for four weeks. Slowly growing colonies with dark-brown to black pigmentation were subcultured in Petri dishes with PDA supplemented with 0.15% malt extract (MPDA) with antibiotics.

Forty blue-stained wood samples (5 × 5 mm), collected adjacent to galleries (eight samples per tree), were surface-sterilized in 70% ethanol for one minute. The samples were then rinsed three times with sterile distilled water, dried on sterile filter paper, and placed on PDA amended with antibiotics. The plates were incubated at 24°C for three weeks. Colonies were selectively subcultured based on growth rate and morphology: fast-growing isolates (>1 cm/day) with cottony aerial mycelium were selected as *Fusarium* candidates, while slow-growing, dark-brown to black colonies were targeted for *Ophiostoma*. All selected isolates were purified on MPDA and grouped into morphotypes based on macroscopic and microscopic characteristics observed using a Jenaval stereomicroscope (Carl Zeiss, Jena).

### 2.3 Molecular identification

The representative isolate F3-1 of the *Fusarium*-like dominant fungal morphotype (DFM), comprising seven isolates from blue-stained wood, and isolate B0922 of the *Ophiostoma*-like DFM, consisting of thirteen isolates from trees and insects, were subjected to molecular identification. Genomic DNA was extracted using the GeneJET Plant Genomic DNA Purification Kit (Thermo Fisher Scientific). The ITS region was amplified for both isolates using primers ITS1F/ITS4R (Gardes & Bruns 1993; White et al. 1990). For the *Ophiostoma*-like isolate B0922, the β-tubulin (*β-TUB*) gene was amplified using primers T10/ Bt2b (O’Donnel and Cigelnik 1997; Glass and Donaldson 1995), and the translation elongation factor 1α (*TEF1-α*) gene region with primers EF1F/EF2R following the protocols of Jacobs et al. (2004). For identification of isolate F3-1, in addition to the ITS region, three loci were amplified: TEF1-α using EF1/EF2 primers (O’Donnell et al. 1998); the DNA-directed RNA polymerase II largest subunit (RPB1) with Fa/ G2R primers (Hofstetter et al. 2007; O’Donnell et al. 2010); and the second largest subunit (RPB2) using 5F2/7Cr primers (Reeb et al. 2004; Liu et al. 1999), following protocols described by Yilmaz et al. (2021). PCR products were purified with QIAquick PCR Purification Kit (Qiagen, Hilden, Germany). Sequencing was carried out by Explogen LLC (EXG, Lviv, Ukraine). Sequences were compared to GenBank via BLASTn, with identification criteria set at >80% coverage, 98–100% similarity for species, and 94–97% for genus. New sequences were deposited in NCBI GenBank.

### 2.4 Phylogenetic analysis

A multilocus sequence analysis (MLSA) of isolate F3-1 was conducted using partial sequences of the *TEF1-α*, *RPB1* and *RPB2* genes. Species identification was supported by comparing these sequences against the FUSARIOID-ID database (Geiser et al. 2004). The top matches were further validated using BLAST on the NCBI platform (http://blast.ncbi.nlm.nih.gov) to assess similarity and infer phylogenetic relationships (Table S1).

Multiple sequence alignments for each locus were generated with the ClustalW algorithm in MEGA 11. The MLSA was performed using MEGA X (v. 11) to construct a Maximum Likelihood (ML) phylogeny based on the Kimura 2-parameter (K2P) model, with branch support evaluated by 1000 bootstrap replicates (Tamura et al. 2021).

### 2.5 Pathogenicity tests

Pathogenicity of fungal isolates was assessed on Scots pine seedlings grown in sandy loam soil under natural irrigation at the Botanical Garden of the National Forestry University of Ukraine (Lviv, Ukraine).

For inoculum preparation, *O. clavatum* isolate B0922 was cultured on 2% malt-extract agar (MEA) at 24° C for 25 days. Agar blocks with fungal mycelium (ca. 5 mm in diameter) were aseptically excised. In May 2023, fifteen three-year-old seedlings were inoculated: bark at 10–12 cm above the root collar was sterilized with 70% ethanol, a 1.5×0.6 cm incision was made to the sapwood, and an agar block with actively growing mycelium was inserted beneath the bark flap, facing the wound. The site was sealed with Parafilm. Controls (15 seedlings) received sterile agar. Seedlings were monitored weekly for 21 weeks, then harvested. Lesion length and depth were measured, and re-isolations were performed from lesion margins on MEA. Fungal identity was confirmed by micromorphology.

Pathogenicity of *Fusarium verticillioides* isolate F3-1 was tested on both seven-day-old and two-year-old seedlings. Seedlings germinated from sterilized seeds were placed on sterile filter paper over 1% water agar in Petri dishes (ten seedlings per treatment). Root zones were inoculated with a spore suspension (7 × 10⁵ CFU/ml) from a ten-day-old PDA culture; sterile water was used as a control. After nine days, seedling health was assessed under a stereomicroscope. Experiments were run in triplicate.

In July 2023, fifteen two-year-old seedlings were inoculated with F3-1. Sites were surface-sterilized as above, and a bark flap was cut to the sapwood. A 2 × 2 mm agar block with actively growing mycelium from a ten-day-old culture was inserted, the flap replaced, and the wound sealed with Parafilm. Controls (15 seedlings) received sterile agar. Seedlings were monitored weekly for eight weeks, then harvested. Necrotic tissue fragments were collected for re-isolation of inoculated fungi.

### 2.6 Antagonism assay

To investigate the interactions between *F. verticillioides* F3-1 and *O. clavatum* В-0922, a dual-culture assay was conducted. Mycelial plugs (5 mm in diameter) were taken from a ten-day-old *F. verticillioides* culture and a fifteen-day-old *Ophiostoma* sp. culture and placed 3 cm apart on the surface of MPDA.

In the control group, mycelial plugs of each fungus were placed individually at the center of separate MPDA plates. Petri dishes were incubated in the dark at 24 °C. Fungal growth was assessed three and seven days after inoculation. All treatments and controls were performed in triplicate.

### 2.7. Statistical analysis

Data are presented as means ± standard errors. The results of the inoculation test were analyzed using a one-way analysis of variance (ANOVA), followed by Dunnett’s test (p ≤ 0.05) to compare each fungal isolate with the control. All analyses were performed using XLSTAT software.

## 3 RESULTS

### 3.1 Morphological and molecular identification of fungal isolates

Twenty-six slow-growing, brown to nearly black isolates resembling *Ophiostoma* were obtained: thirteen from insect abdomens and thirteen from blue-stained sapwood fragments (Figure 1a). Based on morphological characteristics, these were categorized into four distinct morphotypes. The DFM, comprising 13 isolates (7 from insects, 6 from sapwood), was characterized on MPDA by deep dark-brown mycelium with superficial white-beige aerial hyphae and smooth, hyaline margins (Figure 1b), exhibiting an average growth rate of 4.1 ± 0.6 mm/day at 24°C.

**FIGURE 1.**
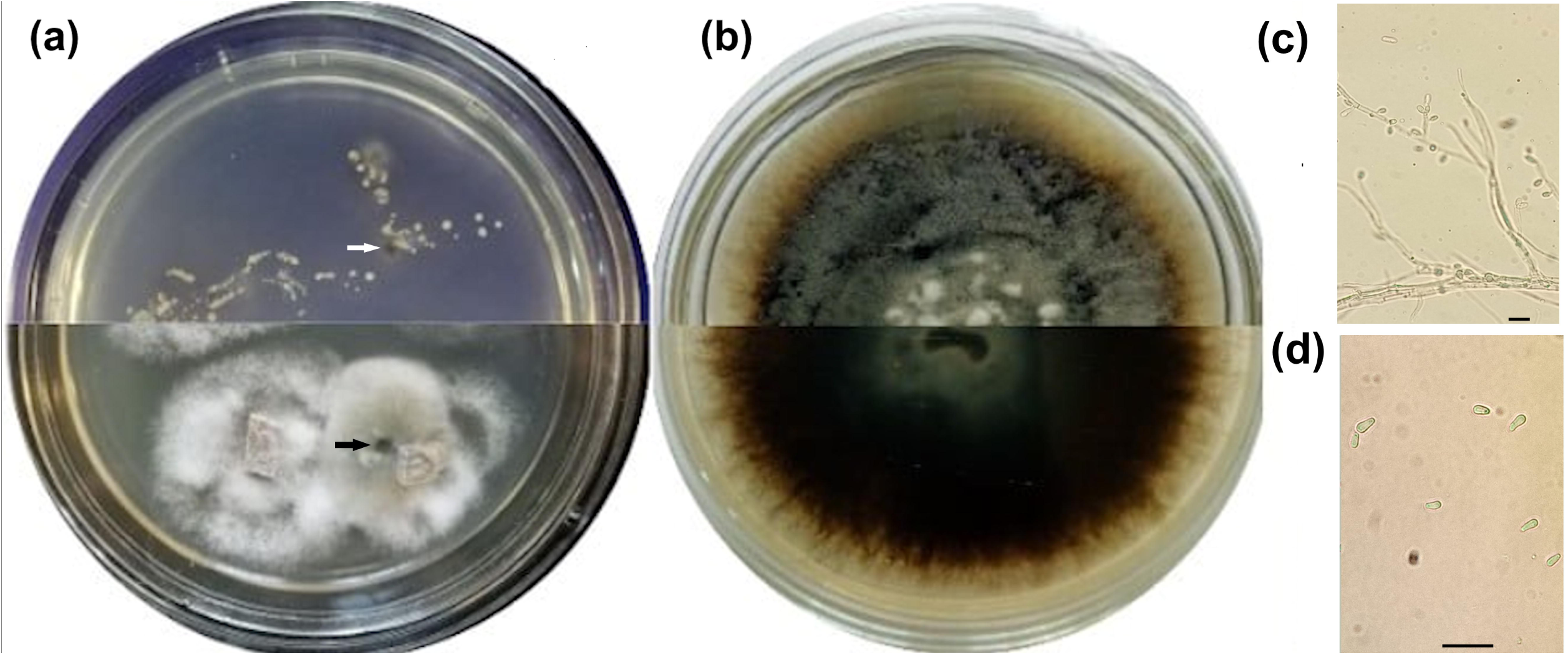
Morphological characteristics of *Ophiostoma clavatum*. (a) Isolates obtained from abdominal segments of *I. acuminatus* (top of plate) and blue-stained sapwood of *P. sylvestris* (bottom); (b) three-week-old culture on MPDA; (c) conidiophore; (d) conidia. Arrows indicate *Ophiostoma*-like colonies. Scale bars: 10 µm

The DFM produced conidiophores typical of the genus *Ophiostoma*, forming small, brush-like clusters (Linnakoski et al., 2016). These structures consisted of thin, elongated, septate hyphae that were often branched, with short lateral branches. Slight swellings indicated the positions of conidiogenous cells. Phialides were small, cylindrical to slightly clavate, usually clustered. Conidia were 3-4 µm long, unicellular, oval to ellipsoidal, borne singly or in short chains (Figures 1c, d). Isolate B0922, a representative of this morphotype, was selected for molecular identification.

The ITS sequence of isolate B0922 (GenBank OR799511.1) showed the highest similarity to *O. clavatum* and *O. brunneolum* (Table 1). While ITS does not fully resolve related taxa, it confirmed affiliation with the *O. clavatum* complex (Linnakoski et al. 2016). For species-level resolution, partial β-TUB and TEF1-α sequences (PX237223.1, PX237224.1) were analyzed, both matching *O. clavatum* strain CMW37983 with the highest identity. Taken together, morphological and molecular data confirm that isolate B0922, obtained from the abdomen of *I. acuminatus*, belongs to *O. clavatum*.

**TABLE 1.** Molecular identification of isolate B0922.

| Isolate | GenBank accession no. | Origin | Gene region | % Sequence coverage | % Identity |
| --- | --- | --- | --- | --- | --- |
| *O. clavatum* isolate CMW37983 | OM501477.1 | Sweden | ITS | 90 | 100 |
| *O. clavatum* isolate CMW37983 | KU094685.1 | Sweden | ITS | 93 | 99.83 |
| *O. brunneolum* isolate CMW44477 | MH144077.1 | China | ITS | 100 | 99.51 |
| *O. hongxingense* strain HXS_66 | MK748194.1 | China | ITS | 100 | 99.06 |
| *O. clavatum* isolate CMW37983 | KU094705.1 | Sweden | β-TUB | 100 | 100 |
| *O. clavatum* isolate CMW41041 | KU094710.1 | Sweden | β-TUB | 100 | 99.69 |
| *O. piliferi* (nom. inval.) strain CXY4047 | OM678560.1 | China | β-TUB | 100 | 96.05 |
| *O. brunneolum* isolate CMW44477 | MH124272.1 | China | β-TUB | 100 | 95.17 |
| *O. clavatum* isolate CMW37983 | KU094759.1 | Sweden | TEF1-α | 85 | 99.72 |
| *O. clavatum* isolate CMW41043 | KU094763.1 | Norway | TEF1-α | 85 | 98.74 |
| *O. brunneolum* isolate CMW44477 | MH124357.1 | China | TEF1-α | 93 | 94.46 |

*Fusarium*-like fungi were detected in 45% of sapwood samples, yielding 18 isolates grouped into six morphotypes. One dominant type, comprising seven isolates from four trees, was chosen for further study.

On MPDA, colonies after 7 days at 24 °C appeared cottony to floccose, with a white to cream margin and violet-gray center (Figure 2a). The reverse side showed yellow to dark-brown pigmentation with concentric rings (Figure 2b). Growth averaged 15.1±1.8 mm/day. Macroconidia (Figure 2c) were slightly falcate, mostly 3-septate, 20–40 µm long. Microconidia (Figure 2d) were unicellular, hyaline, oval to club-shaped, 5–8 µm, produced in chains or false heads, traits typical of the *Fusarium fujikuroi* species complex (FFSC) (Bao et al. 2023).

**FIGURE 2.**
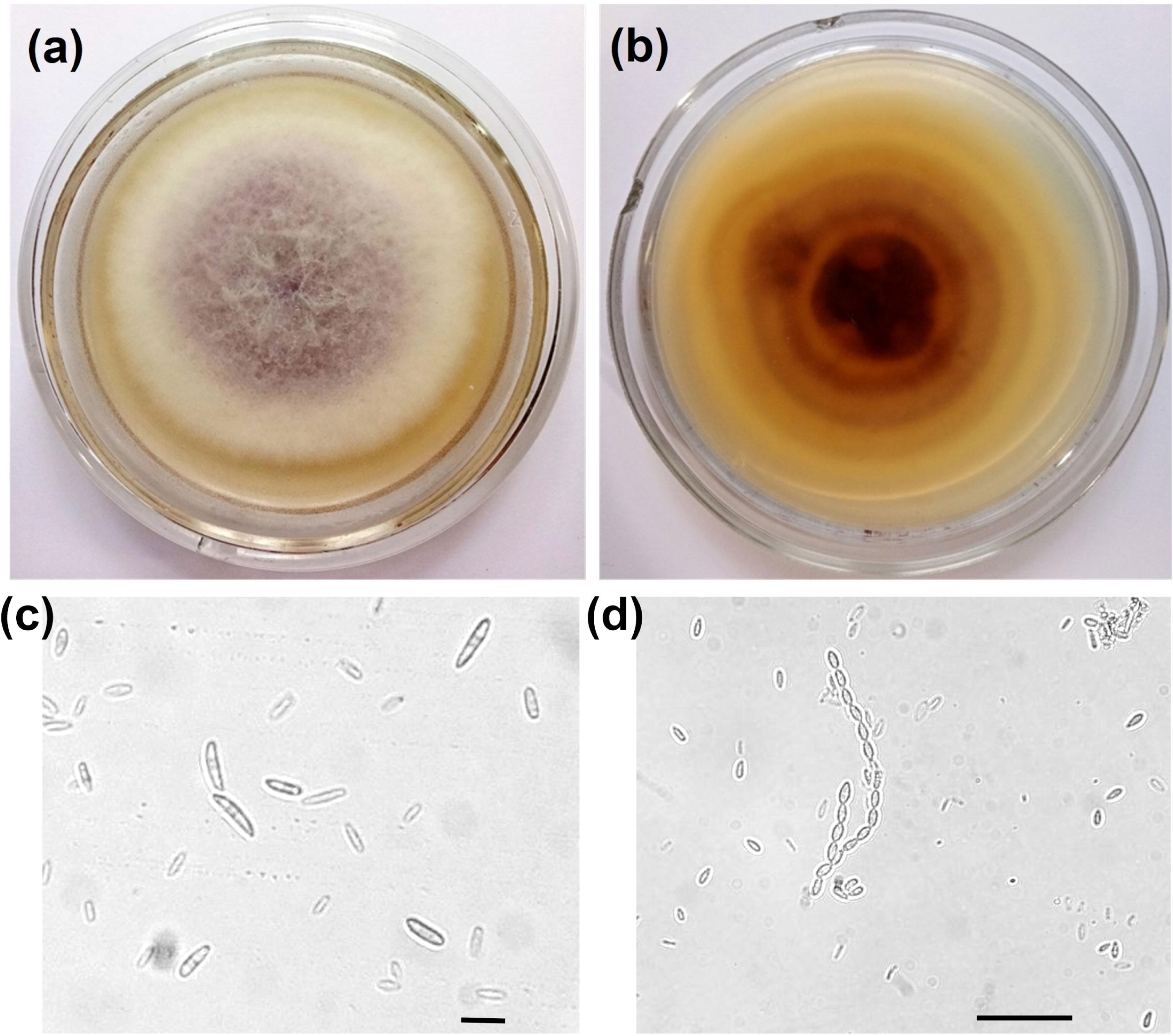
Morphological characteristics of *Fusarium verticillioides*. (a, b) Seven-day-old culture on MPDA (surface and reverse); (c) macroconidia; (d) microconidia. Scale bars: 20 µm.

ITS sequencing of isolate F3-1 (PQ488808), representative of this morphotype, showed 99.64% identity (98% coverage) to *Fusarium cf. fujikuroi* isolate 56339C (PP804370.1) and *F. verticillioides* isolate BFO (PP434657.1), and 99.45% identity to *F. circinatum*, *F. coicis*, *F. temperatum*, and *F. napiforme*. All belong to the FFSC.

The ML phylogenetic tree, constructed from the combined sequence data of three loci (tef1-α, rpb1, and rpb2), clearly demonstrates two major clades with strong statistical support (bootstrap = 100). The first clade comprised diverse *Fusarium* species, including *F. agapanthi*, *F. ananatum*, *F. globosum*, *F. bactridioides*, *F. circinatum*, *F. begoniae*, and *F. dlaminii*. The second major clade contained representatives of *F. verticillioides* together with *F. concentricum*, *F. annulatum*, and *F. xylarioides* (Figure 3). Isolate F3-1 was placed within the *F. verticillioides* group and clustered most closely with strain LC2818, supported by a bootstrap value of 90. This confirms the identity of isolate F3-1 as *F. verticillioides*.

**FIGURE 3.**
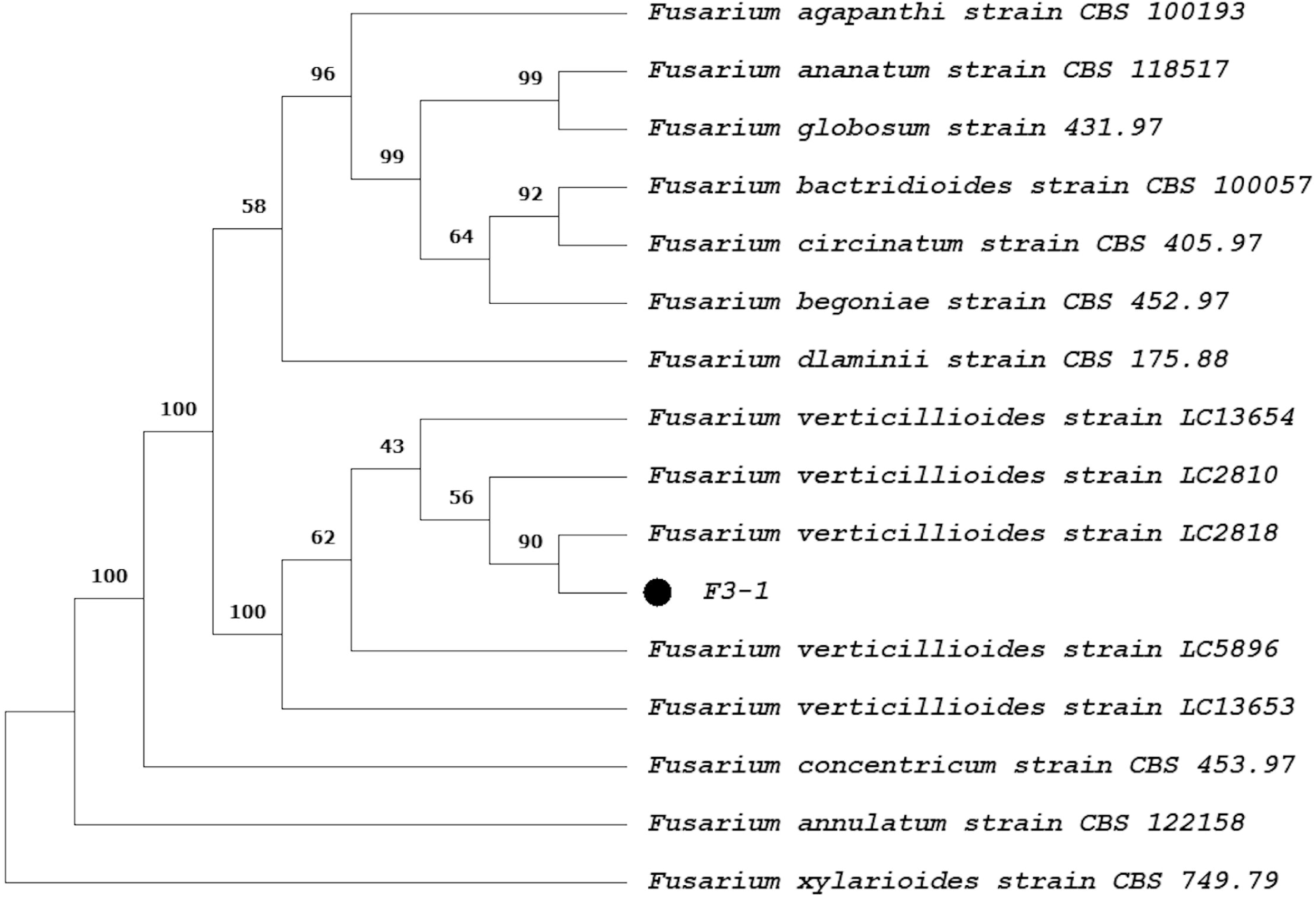
Maximum likelihood (ML) tree of *F. verticillioides* isolate F3-1 based on a combined three-gene dataset (*tef1-α*, *rpb1*, and *rpb2*) generated in MEGA 11.0. Values above nodes represent bootstrap support (1000 replicates)

### 3.2 Pathogenicity tests

Twenty-one weeks after inoculation, 60% of *P. sylvestris* seedlings inoculated with *O. clavatum* developed moderate bleeding lesions on the outer bark (Figure 4a); the remainder showed only slight exudation. Control seedlings exhibited dry wounds with localized discoloration only. Inoculated plants formed elongated dark-brown lesions extending vertically from the inoculation point (Figure 4b). Lesions in controls averaged 15.2 ± 1.3 mm long and 1.2 ± 0.1 mm deep, whereas in inoculated seedlings reached 30.5 ± 3.6 mm and 4.4 ± 0.6 mm, respectively. Differences were statistically significant (p < 0.001). The fungus was reisolated from 100% of inoculated seedlings and matched the original inoculum morphologically (Figure 4c). No fungi were reisolated from control plants.

**FIGURE 4.**
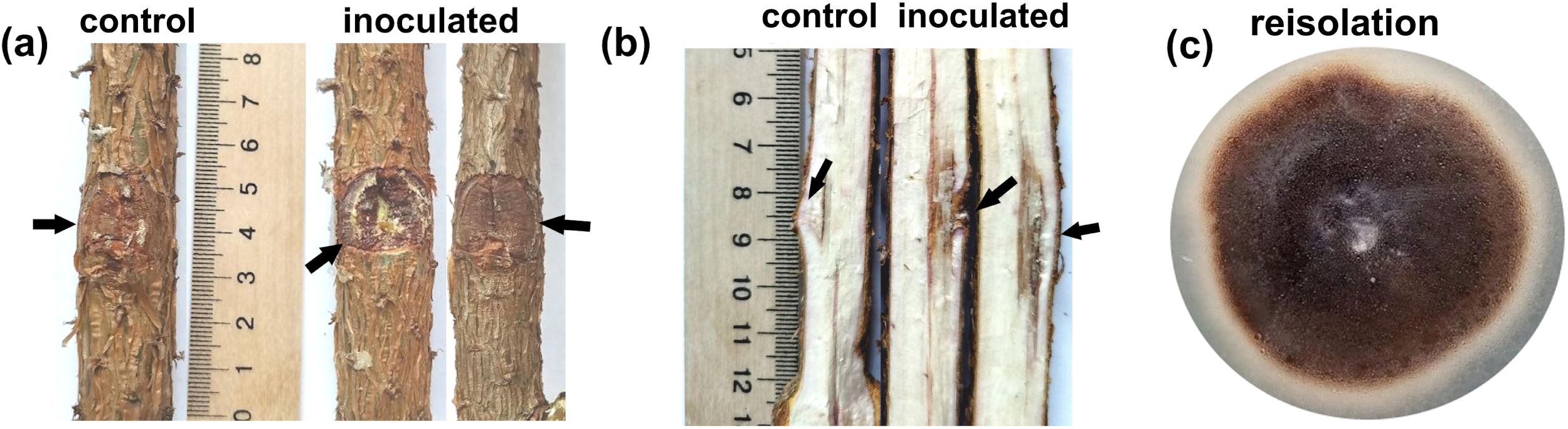
Pathogenicity of *O. clavatum*. Three-year-old *P. sylvestris* seedlings were inoculated beneath bark flaps with agar blocks containing *O. clavatum* mycelium (inoculated) or sterile agar (control). Symptoms after 21 weeks: (a) inoculation site on stem; (b) longitudinal stem section through the inoculation site; (c) culture reisolated from inoculated seedlings. Arrows indicate inoculation points

Inoculation of seven-day-old seedlings with *F. verticillioides* caused symptoms in 84.3 ± 4.6% of plants, including root and hypocotyl browning, white aerial mycelium on hypocotyls, black necrotic strands along vascular tissues, tissue maceration, and cotyledon damage. Controls showed no symptoms (Figure 5).

**FIGURE 5.**
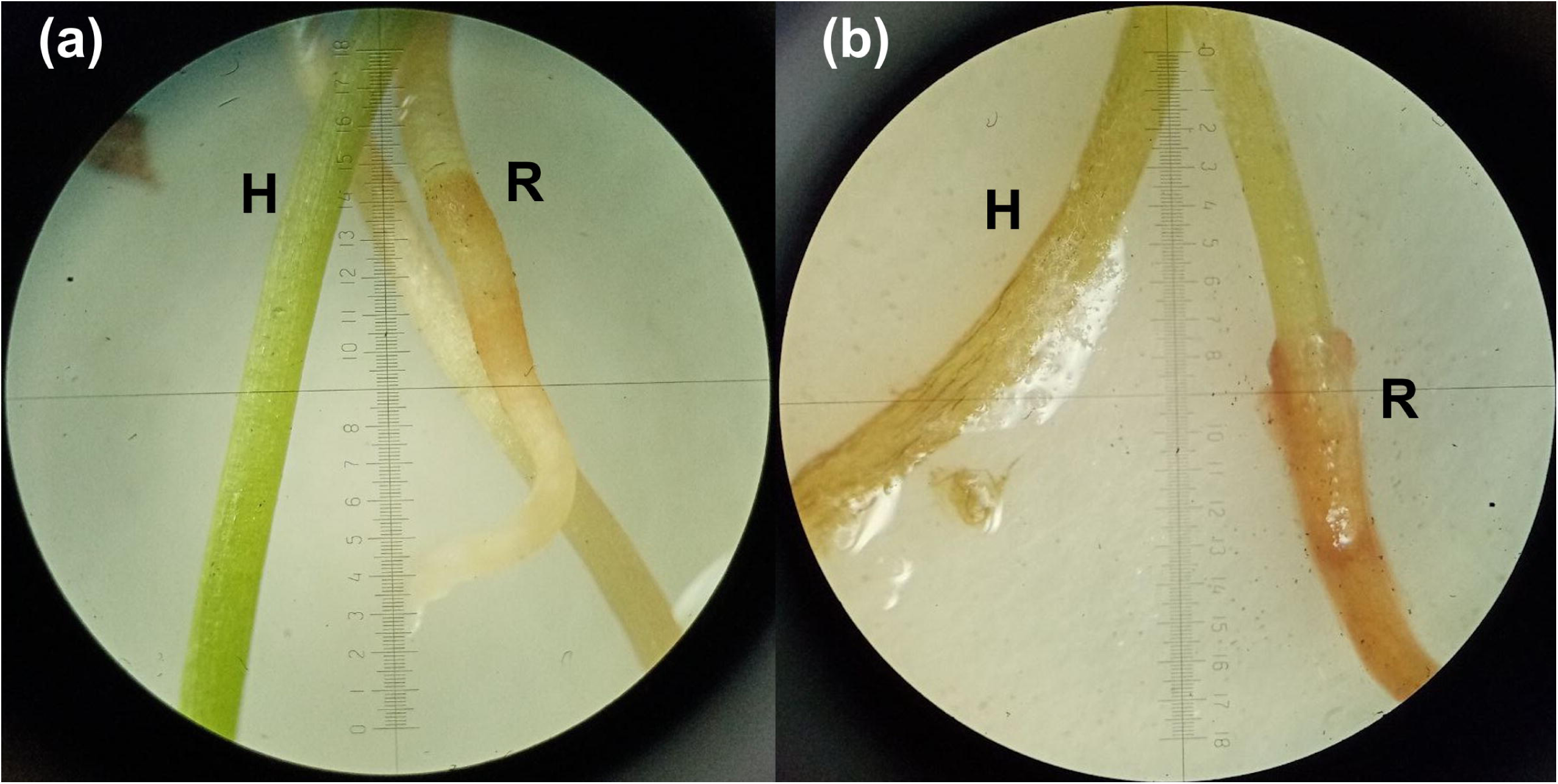
Pathogenicity test of *F. verticillioides* on seven-day-old *P. sylvestris* seedlings. (a) Control seedlings showed no pathological changes; (b) inoculated seedlings displayed root and hypocotyl browning, white aerial mycelium on hypocotyls, and tissue maceration. Abbreviations: H – hypocotyl, R – root.

In two-year-old seedlings inoculated beneath bark flaps, all plants developed disease. Resin exudation occurred in 73.3% of seedlings, and some showed unhealed wounds (Figure 6a). Controls exhibited complete wound closure by cambial growth. Necrotic lesions extended 5.8 ± 0.8 mm from the incision, with cross-sections showing wood discoloration absent in controls (Figure 6b). Necrosis reached the second annual ring in 60.4% of seedlings and the first ring in the remainder. The pathogen was reisolated from all inoculated plants.

**FUGURE 6.**
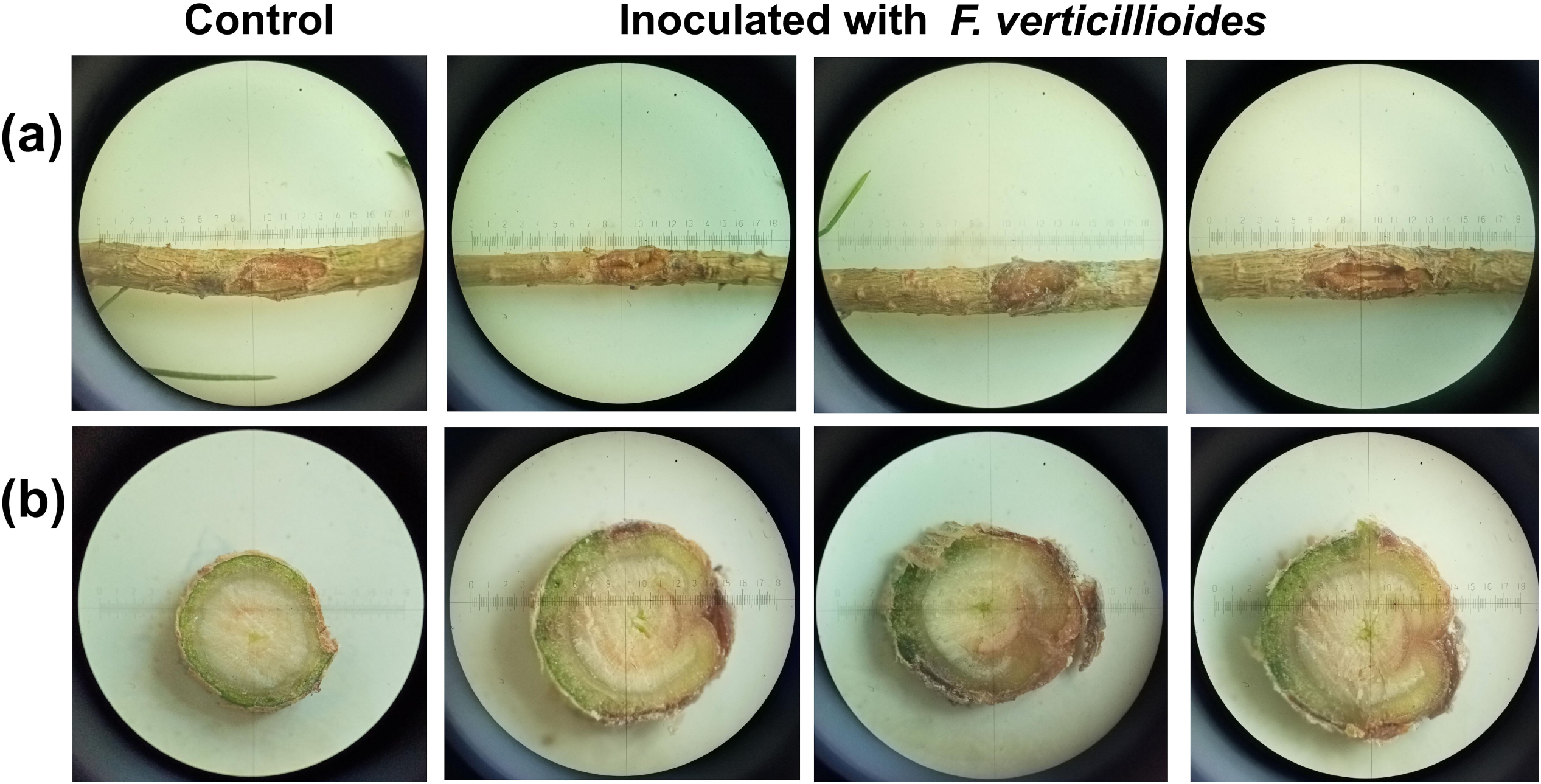
Pathogenicity test of *F. verticillioides* on two-year-old seedlings. Representative symptoms after 8 weeks of incubation on the stem in the inoculation site (a) and on cross-section through the inoculation site (b)

### 3.3 Antagonism assay

After three days in dual culture, *F. verticillioides* showed rapid mycelial expansion with diffuse pink pigmentation at the colony center. *O. clavatum* formed colonies averaging 14.1 ± 1.2 mm, comparable to its control, but developed a dense structure with a darkened center and mycelial growth directed away from the co-inoculant (Figure 7a).

**FIGURE 7.**
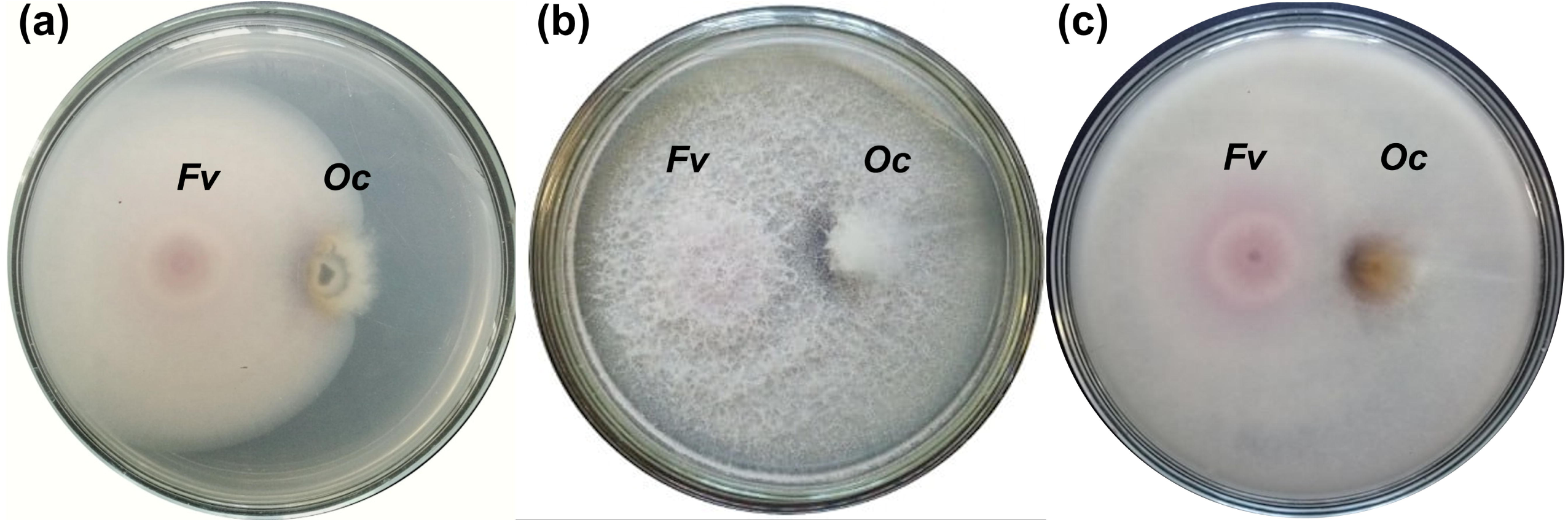
Dual-culture assay of *F. verticillioides* F3-1 (*Fv*) and *O. clavatum* B0922 (*Oc*) on MPDA plates. (a) Reverse view after three days; (b) front view after seven days; (c) reverse view after seven days

By day seven, *F. verticillioides* had colonized most of the plate, producing dense, cottony white mycelium on the surface and distinct pink pigmentation on the reverse. In contrast, *O. clavatum* formed smaller, compact colonies with dense aerial mycelium and a dark-brown zone at the inoculation site (Figure 7b, c). Growth of *O. clavatum* was restricted compared with controls. Colony size in dual culture averaged 16.1 ± 1.2 mm versus 29.4 ± 1.6 mm in control, confirming the strong suppressive effect of *F. verticillioides*.

## 4 DISCUSSION

This study provides the first report of *O. clavatum* associated with *I. acuminatus* in Scots pine forests of western Ukraine. Molecular identification of isolate B0922 (ITS, TUB-2, TEF1-α) showed 99.72–100% identity with *O. clavatum* strain CMW 37983 from *I. acuminatus* infesting *P. sylvestris* in Sweden. This widespread beetle–fungus association has been documented in Austria, France, Germany, Norway, Sweden, and the former Yugoslavia (Francke-Grosmann, 1952, 1963; Lieutier et al. 1991; Linnakoski et al. 2016; Waalberg, 2015), but until now not in Ukraine or Poland (Davydenko et al. 2017; Davydenko and Baturkin, 2020; Jankowiak et al. 2023). Its presence in western Ukraine suggests the association may be distributed throughout Europe.

Fungal diversity in *I. acuminatus* varies with isolation method (Kirisits, 2004; Linnakoski et al. 2016; Papek et al. 2024). Beetle exoskeletons and galleries accumulate spores from trees, mites, and other beetles, typically harboring diverse communities where taxa of interest may be suppressed in culture by faster-growing fungi (Furniss et al. 1990). In contrast, the gut hosts a smaller, more stable core mycobiome likely important for beetle biology (Hu et al. 2015; Isb et al. 2015). *Ophiostoma* species are frequently reported from the guts of pine beetles, including *I. acuminatus* and *I. sexdentatus* (Chakraborty et al. 2020). In this study, *Ophiostoma*-like isolates representing four morphotypes were obtained from all insect samples, with *O. clavatum* dominant in both beetles and blue-stained sapwood samples.

Inoculation of three-year-old Scots pine seedlings with *O. clavatum* caused moderate lesions in all plants. In a comparable study under less stressful conditions, only 45% of seedlings were infected (Isberg et al. 2022). The higher severity of seedlings in our study may reflect a longer trial, larger wounds, higher inoculum, and environmental stress. Nevertheless, mortality was absent, confirming that *O. clavatum* is pathogenic yet weakly virulent (Guérard et al. 2000; Villari et al. 2012; Isberg et al. 2022). Such fungi may instead weaken host defenses, facilitating beetle colonization (Guérard et al. 2007; Villari et al. 2012).

The dominant *Fusarium* morphotype from blue-stained wood was identified by TEF1-α, RPB1, and RPB2 sequences as *F. verticillioides*, a member of the *F. fujikuroi* species complex. Globally, *F. verticillioides* is a major maize pathogen causing ear rot (Deepa and Sreenivasa, 2017), but it also affects conifers, causing damping-off in several pine species (Olaizola et al. 2023) and resinous canker in *Pinus greggii* (De León-Torres et al. 2023). Isolate F3-1 proved pathogenic on pine seedlings, producing cortex disintegration, resinous lesions, and tissue necrosis. Its ability to damage two-year-old seedlings underscores the need to examine its role in mature pine sapwood. Like ophiostomatoid fungi, it may disrupt water transport or degrade tissues to aid beetle development, thereby intensifying pest–pathogen interactions, as seen in the *F. solani*– *Hypothenemus hampei* symbiosis (Morales-Ramos et al. 2000).

The detection of *F. verticillioides* in the xylem raises questions about its entry pathways. Potential routes include beetle-mediated transmission (either externally or via the gut), colonization through insect feeding galleries, or environmental entry via the roots. The latter hypothesis is supported by its recent identification as an endophyte in *P. sylvestris* var. *mongolica* (Ren et al., 2024).

In the dual-culture assay, *F. verticillioides* rapidly colonized the plates and suppressed the growth of *O. clavatum*. The latter responded with localized pigmentation, indicating stress or an antagonistic interaction. This *in vitro* result suggests that their interaction *in situ* is likely competitive rather than neutral. Consequently, the superior competitiveness of *F. verticillioides* may restrict the colonization of *O. clavatum* in sapwood by suppressing its growth and outcompeting it for resources. However, *in vivo* outcomes may differ from the dual-culture results due to the structural and nutritional complexity of wood and the influence of other microbial associates.

## CONCLUSIONS

This study provides the first evidence of *O. clavatum* associated with *Ips acuminatus* in Ukraine, expanding its known range in Europe. Inoculation trials confirmed that *O. clavatum* is phytopathogenic but weakly virulent; it likely contributes to host weakening rather than direct mortality. In contrast, *F. verticillioides* demonstrated strong pathogenicity on Scots pine seedlings and competitive dominance over *O. clavatum*, suggesting a possible involvement in pine decline. These findings highlight the need for future studies to elucidate the multipartite interactions within bark beetle-vectored fungal communities, which are important for understanding their role in tree susceptibility and forest decline dynamics.

## ACKNOWLEDGMENTS

The study was funded by the National Research Foundation of Ukraine (project 2021.01/0184). This study was partially funded by “Presidential Discretionary-Ukraine Support Grants” SFI-PD-Ukraine-00014576 from Simons Foundation, [VAK]

## AUTHOR CONTRIBUTIONS

Conceptualization, [VAK, YMY]; methodology, [VAK, YMY, OYK, VOK]; validation, [HTK, VKZ]; formal analysis, [OYK, MMK, YIS]; investigation, [YMY, OYK, VAK]; resources, [VKZ, VVL, HTK, VOK]; data curation, [VAK, OYK, MMK]; writing – original draft preparation, [VAK]; writing – review and editing, [VAK, YIS]; visualization, [YMY, OYK]; supervision, [VAK]; project administration, [VAK, HTK]; funding acquisition, [YMY, OYK, VOK, YIS, MMK, VVL, HTK, VAK]. All authors have read and agreed to the published version of the manuscript.

