## Supplementary material for "Pathogenic Fusarium verticillioides and Ophiostoma clavatum Associated with Ips acuminatus in Ukraine": Table S1

**Table S1.** Strains used for Maximum Likelihood (ML) phylogenetic analysis, including their source metadata and percent identity to isolate F3-1

| **Species** | **Origin** | **Source** | ***TEF1-α*** | | ***RPB1*** | | ***RPB2*** | |
| --- | --- | --- | --- | --- | --- | --- | --- | --- |
|  |  |  | **GenBank ID** | **Identity,**  **%** | **GenBank**  **ID** | **Identity,**  **%** | **GenBank**  **ID** | **Identity,**  **%** |
| *Fusarium agapanthi* strain *CBS 100193* | New Zealand | *Agapanthus praecox* | MW401959 | 90.59 | MW402491 | 97.27 | MW402727 | 95.52 |
| *Fusarium ananatum* strain *CBS 118517* | South Africa | *Ananas comosus* | MN533988 | 92.00 | MW402508 | 96.54 | MN534229 | 95.39 |
| *Fusarium annulatum* strain *CBS 115.97* | Italy | *Dianthus caryophyllus* | MW401973 | 90.39 | MW402503 | 95.81 | MW402785 | 95.76 |
| *Fusarium bactridioides* strain *CBS 100057* | USA | *Cronartium conigenum* on *Pinus leiophylla* | MN533993 | 90.62 | MW402490 | 95.81 | MN534235 | 94.33 |
| *Fusarium begoniae* strain *CBS 452.97* | Germany | *Begonia elatior* hybrid | MN533994 | 91.83 | MW402675 | 96.90 | MN534243 | 95.27 |
| *Fusarium circinatum* strain *CBS 405.97* | USA | *Pinus radiata* | MN533997 | 91.59 | MW402656 | 96.17 | MN534252 | 93.97 |
| *Fusarium dlaminii* strain *CBS 175.88* | South Africa | *Zea mays* soil | MN534002 | 92.30 | MW402623 | 97.45 | MN534256 | 95.39 |
| *Fusarium globosum* strain 431.97 | South Africa | *Zea mays* seed | MW402131 | 90.39 | MW402669 | 96.17 | MW402816 | 95.98 |
| *Fusarium xylarioides* strain *CBS 749.79* | Guinea | *Coffea canephora* | MN534049 | 91.87 | MW402702 | 98.18 | MN534259 | 95.39 |
| *Fusarium concentricum* strain *CBS 453.97* | Guatemala | *Musa sapientum* | MN533998 | 91.42 | MW402676 | 95.45 | MN534264 | 95.39 |
| *Fusarium verticillioides* strain *LC13654* | USA | *Glycine max* | [MW580505](javascript:void(0)) | 100 | [MW024493](javascript:void(0)) | 99.64 | [MW474451](javascript:void(0)) | 99.66 |
| *Fusarium verticillioides* strain *LC2810* | China | Bamboo | [MW580507](javascript:void(0)) | 100 | [MW024495](javascript:void(0)) | 99.64 | [MW474453](javascript:void(0)) | 99.66 |
| *Fusarium verticillioides* strain *LC2818* | China | *Physosfegia virginiana* | [MW580508](javascript:void(0)) | 100 | [MW024496](javascript:void(0)) | 99.64 | [MW474454](javascript:void(0)) | 99.66 |
| *Fusarium verticillioides* strain *LC5896* | China | Submerged wood | [MW580509](javascript:void(0)) | 100 | [MW024497](javascript:void(0)) | 99.64 | [MW474455](javascript:void(0)) | 99.66 |
| *Fusarium verticillioides* strain *LC13653* | Brazil | *Glycine max* | [MW580504](javascript:void(0)) | 100 | [MW024492](javascript:void(0)) | 99.64 | [MW474450](javascript:void(0)) | 99.43 |
| *Fusarium verticillioides* isolate *F3-1* | Ukraine | *Pinus sylvestris* | PQ488808 |  | PV976806 |  | PV976805 |  |
